# Substrate displacement of CK1 C-termini regulates kinase specificity

**DOI:** 10.1101/2023.06.30.547285

**Authors:** Sierra N. Cullati, Kazutoshi Akizuki, Jun-Song Chen, Kathleen L. Gould

**Author notes:** **Teaser:** We propose a model for CK1 substrate specificity based on autophosphorylation state.

## Abstract

CK1 kinases participate in many signaling pathways; how these enzymes are regulated is therefore of significant biological consequence. CK1s autophosphorylate their C-terminal non-catalytic tails, and eliminating these modifications increases substrate phosphorylation in vitro, suggesting that the autophosphorylated C-termini act as inhibitory pseudosubstrates. To test this prediction, we comprehensively identified the autophosphorylation sites on *Schizosaccharomyces pombe* Hhp1 and human CK1ε. Peptides corresponding to the C-termini interacted with the kinase domains only when phosphorylated, and phosphoablating mutations increased Hhp1 and CK1ε activity towards substrates. Interestingly, substrates competitively inhibited binding of the autophosphorylated tails to the substrate binding grooves. The presence or absence of tail autophosphorylation influenced the catalytic efficiency with which CK1s targeted different substrates, indicating that tails contribute to substrate specificity. Combining this mechanism with autophosphorylation of the T220 site in the catalytic domain, we propose a displacement specificity model to describe how autophosphorylation regulates substrate specificity for the CK1 family.

## Introduction

CK1 enzymes are conserved, ubiquitous kinases that regulate essential cellular pathways including Wnt signaling, cell division, endocytosis, circadian rhythms, and DNA repair (Cheong and Virshup, 2011; Knippschild et al., 2014). Like other multifunctional kinases, CK1s must be regulated in space and time in order to specifically target their substrates in each of the pathways they participate in. Because CK1s prefer substrate motifs that have been previously phosphorylated (Flotow et al., 1990), the action of substrate-priming kinases, or the opposing phosphatases, is considered a key mechanism of achieving CK1 substrate specificity (Bhandari et al., 2013; Ishiguro et al., 2010; Rivers et al., 1998; Zeng et al., 2005). However, several important substrates (e.g. PER2 in circadian rhythms (Narasimamurthy et al., 2018), LRP6 in Wnt signaling (Swiatek et al., 2006), Sid4 in a mitotic checkpoint (Johnson et al., 2013), Rec11 in meiotic recombination (Phadnis et al., 2015; Sakuno and Watanabe, 2015)) are not primed by other kinases, indicating that there must exist other mechanisms to control CK1 activity.

CK1 family members have related catalytic domains (53%-98% sequence identity), a conserved extension to that kinase domain (KDE) that is important for enzyme stability and activity (Cheong and Virshup, 2011; Elmore et al., 2018; Greer and Rubin, 2011; Knippschild et al., 2014; Ye et al., 2016), and divergent C-terminal tails. In addition to catalytic domain autophosphorylation that occurs in many CK1 enzymes (Cullati et al., 2022), all C-terminal tails appear to serve as substrates of autophosphorylation, and the phosphorylated C-termini are proposed to inhibit enzyme activity by acting as pseudosubstrates (Cegielska et al., 1998; Gietzen and Virshup, 1999; Graves and Roach, 1995; Hoekstra et al., 1994).

While there is abundant evidence that CK1 enzymes autophosphorylate their C-termini, and truncation or dephosphorylation of the tail increases kinase activity in vitro (Cegielska et al., 1998; Gietzen and Virshup, 1999; Graves and Roach, 1995; Hoekstra et al., 1994), an interaction between the phosphorylated tail and the kinase domain has not been demonstrated, and questions remain about how this regulatory mechanism functions. It is unclear whether truncation of the tail and dephosphorylation have equivalent effects on the enzyme. The affinity with which this interaction may occur in the context of substrate binding, and therefore how the plethora of CK1 substrates are regulated by autophosphorylation, is only beginning to be understood. While some autophosphorylation sites have been identified in CK1ε (Gietzen and Virshup, 1999), the full complement of autophosphorylation sites, their cellular functions, and how autophosphorylation may affect different substrates has not been determined for any CK1 enzyme.

In order to investigate how CK1 family proteins are regulated by their C-termini, we focused on human CK1ε and *Schizosaccharomyces pombe* Hhp1 as representative CK1 enzymes. Hhp1 is one of two soluble CK1 enzymes in *S. pombe* (Hhp2 is the other), and these two yeast enzymes are highly related to CK1δ and CK1ε in human (Dhillon and Hoekstra, 1994; Hoekstra et al., 1994). Autophosphorylation of all four of these enzymes has been observed previously (Cegielska et al., 1998; Cullati et al., 2022; Gietzen and Virshup, 1999; Graves and Roach, 1995; Hoekstra et al., 1994). Here, we identified all residues in the Hhp1 and CKε C-termini that are autophosphorylated in vitro; these sites are also targeted in vivo. Preventing phosphorylation of these specific sites increased kinase activity toward substrates. Reciprocally, the respective C-terminal phosphopeptides bound to their kinase domains, and this binding competitively inhibited substrate phosphorylation. We confirmed that autophosphorylated C-terminal peptides and substrates interacted with the kinase domain via the same positively charged amino acids in the binding groove. Critically, we found that substrates have a much greater affinity for the substrate binding groove than the phosphorylated C-termini, and that competition between the autophosphorylated C-termini and substrates for access to the kinase domains regulated the efficiency with which different substrates were targeted. This supports an updated paradigm for the regulation of CK1 family kinases via autophosphorylation, in which the phosphostate of CK1 can influence which substrates have high affinity for the catalytic domain.

## Results

### Hhp1 autophosphorylates six sites on its C-terminus

CK1 enzymes phosphorylate their C-terminal, non-catalytic tails (Cegielska et al., 1998; Gietzen and Virshup, 1999; Graves and Roach, 1995; Hoekstra et al., 1994), but the full complement of autophosphorylation sites has not been identified for any one enzyme. In order to identify autophosphorylation sites on *S. pombe* Hhp1, we analyzed the recombinant kinase by mass spectrometry (Figure S1A). Many of the phosphopeptides that we identified contained multiple serines and threonines, leading to ambiguous localization of phosphorylation sites. Therefore, we eliminated false positives and verified candidate sites by generating alanine substitution mutants at each candidate site and performing phosphopeptide mapping of the autophosphorylated, recombinant proteins (Figure S1B). In total, we found that six sites in the Hhp1 C-terminus can be autophosphorylated: Ser326, Ser327, Thr345, Thr346, Ser354, and Thr356 (Figure 1A). Phosphorylated Ser354 and Thr356 have also been observed in large-scale phosphoproteomics experiments (Carpy et al., 2014; Kettenbach et al., 2015; Koch et al., 2011; Swaffer et al., 2018). A mutant with all six sites mutated to alanine (Hhp1-6A) produces an autophosphorylation map identical to that of C-terminally truncated Hhp1 (Hhp1ΔC, amino acids 1-296) (Figure S1B). The remaining phosphopeptides are due to T221 and T222 autophosphorylation of the kinase domain (Cullati et al., 2022). Not only could CK1 autophosphorylate the C-terminal sites in the full-length protein, but we expressed and purified amino acids 297-365 of Hhp1 (Cter) fused to GST, and Hhp1ΔC was able to utilize wildtype Cter, but not Cter-6A, as a substrate *in trans* (Figure 1B). These data support the conclusion that all Hhp1 C-terminal sites that can be autophosphorylated in vitro were identified.

**Figure 1:**
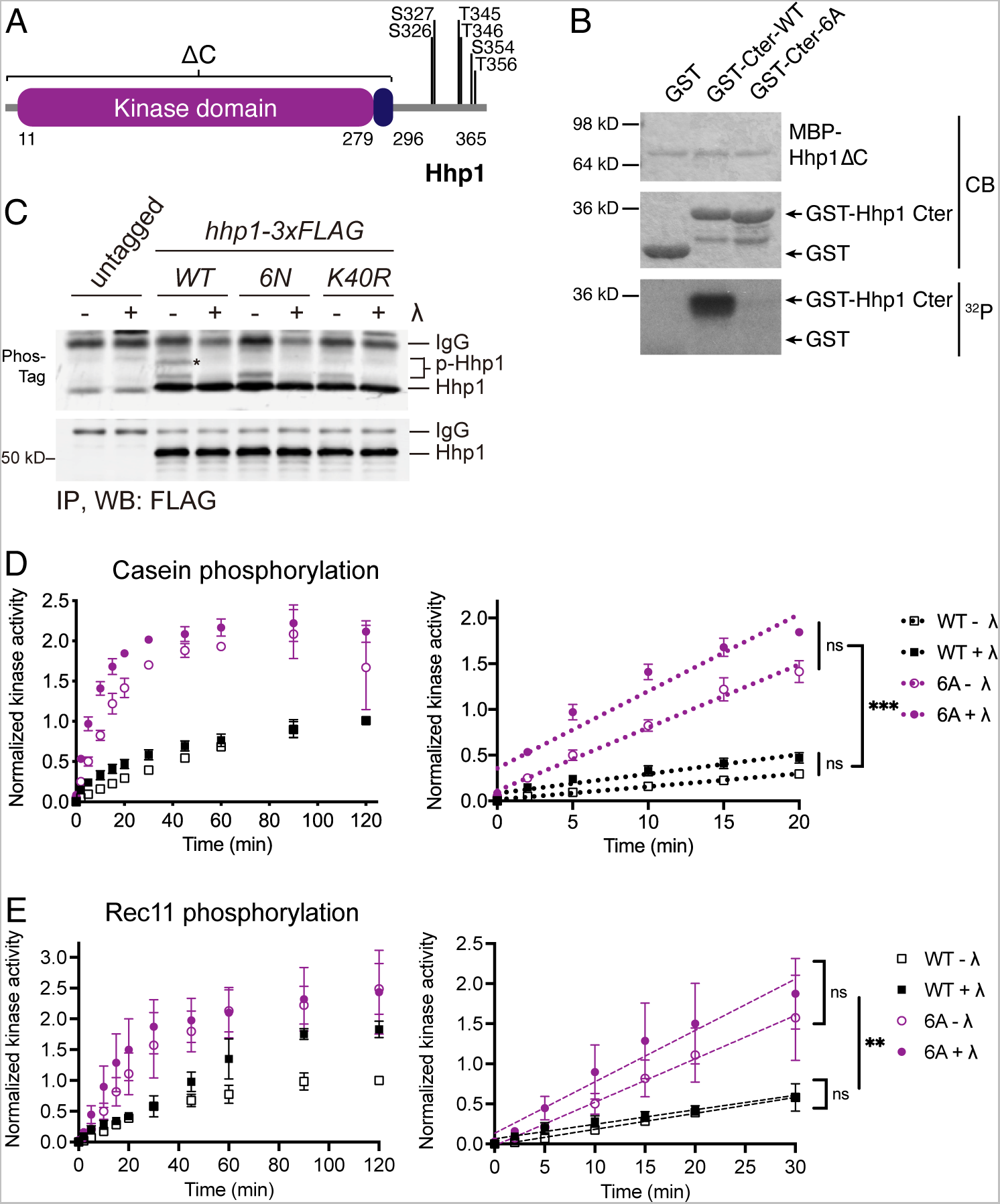
Autophosphorylation at six sites inhibits Hhp1 activity. (A) Domain structure of *S. pombe* Hhp1 showing autophosphorylation sites identified by mass spectrometry and confirmed by phosphopeptide mapping. (B) MBP-Hhp1ΔC was incubated with GST-Hhp1-Cter and γ-[^32^P]-ATP at 30°C for 2 h. Phosphorylation was detected by autoradiography (^32^P) and total protein by Coomassie (CB). (C) *hhp1-wt, hhp1-6N,* and *hhp1-K40R* were tagged with 3xFLAG and the resultant proteins were immunoprecipitated from denatured lysates with anti-FLAG antibody. IPs were treated +/- lambda phosphatase (λ) and electrophoretic mobility shifts on 6% gels containing 20 µM Phos-Tag were detected with anti-FLAG antibody (upper). Total Hhp1-3xFLAG was visualized on gels without Phos-Tag (lower). Asterisk (*) indicates autophosphorylated Hhp1-3xFLAG. (D-E) MBP-Hhp1-WT or -6A was treated +/- λ, then incubated with substrate and γ-[^32^P]-ATP at 30°C. Reactions were quenched at timepoints from 0-120 min, and casein phosphorylation (D) or Rec11 phosphorylation (E) was measured on a phosphorimager. Data from three independent replicates is shown as the mean ± SD. The full timecourse is shown on the left; the initial rate was determined by the slope of the linear section of the curve shown on the right. ** = p < 0.001, *** = p < 0.0002, ns = not significant by one-way ANOVA of slopes.

One advantage of studying mechanisms regulating CK1 enzymes using Hhp1 as a model is that the effects of mutations ablating autophosphorylation can be easily tested in vivo. We had previously found that C-terminally truncating the enzyme, which would have removed all C-terminal autophosphorylation sites, had no discernable effect on cell growth or subcellular localization of Hhp1 (Elmore et al., 2018). We anticipated the same for the *hhp1-6A* mutant, and to test this, we integrated *hhp1-6A* at the endogenous locus. We found that *hhp1-6A* but not *hhp1-6N*, which also eliminates C-terminal autophosphorylation by substituting with asparagine, had a growth defect, indicating a partial loss of function (Figure S2A). Loss of function was confirmed in a *hhp1^+^/hhp1-6A* diploid (Figure S2B). We surmise that the more polar amino acid preserves function better than alanine; therefore, we used *hhp1-6N* for our in vivo studies. Hhp1-6N tagged with mNeonGreen (mNG) localized indistinguishably from Hhp1-mNG and was present in the cytoplasm and nucleus, and also concentrated at the spindle pole body and cell division site (Figure S2C) (Elmore et al., 2018). To determine whether the C-terminal autophosphorylation sites identified in vitro were also targeted in vivo, we tagged *hhp1, hhp1-6N,* and kinase-dead *hhp1-K40R* at their endogenous loci with 3xFLAG and examined their electrophoretic mobility. Immunoprecipitating and treating with lambda phosphatase collapsed Hhp1 WT-3xFLAG on a gel containing Phos-Tag reagent (Figure 1C). Conversely, Hhp1-6N-3xFLAG and Hhp1-K40R-3xFLAG migrated faster than WT and identically to each other (Figure 1C), indicating that these sites are autophosphorylated in vivo. Both Hhp1-6N-3xFLAG and Hhp1-K40R-3xFLAG lost the major phosphorylated species present in Hhp1 WT-3xFLAG while retaining a minor phosphorylated species, suggesting that Hhp1 can also be phosphorylated by other cellular kinases.

Truncation or phosphatase treatment of CK1δ, CK1ε, Hhp1, and Hhp2 increases phosphorylation of the model substrate casein (Cegielska et al., 1998; Cullati et al., 2022; Gietzen and Virshup, 1999; Graves and Roach, 1995; Hoekstra et al., 1994). This predicts that preventing autophosphorylation of the Hhp1 C-terminal sites would similarly increase Hhp1’s ability to phosphorylate casein and other substrates. We measured the rate of casein phosphorylation by recombinant Hhp1-6A and Hhp1-6N, obtaining identical results with these two mutants in vitro (Figure 1D and Figure S2D). Consistent with the literature, Hhp1-6A and Hhp1-6N were about two-fold more active than the wildtype enzyme, and the initial velocities of substrate phosphorylation by Hhp1-6A and Hhp1-6N were significantly greater than that of Hhp1-WT (Figure 1D). These results were recapitulated using the N-terminus of Rec11 (GST-Rec11-N3, consisting of amino acids 1-33; Sakuno and Watanabe, 2015), a physiological Hhp1 substrate involved in meiosis (Phadnis et al., 2015; Sakuno and Watanabe, 2015) (Figure 1E and Figure S2E). Because Hhp1-6A and Hhp1-6N cannot become autophosphorylated on their C-termini during the course of the experiment, these mutants were even more active than dephosphorylated wildtype Hhp1 after two hours (Figure 1D-E and Figure S2D-E).

### The C-terminus of Hhp1 interacts with the kinase domain in a phosphorylation-dependent manner

The idea that autophosphorylated CK1 C-termini can act as pseudosubstrates predicts that they interact with their kinase domains (Cegielska et al., 1998; Gietzen and Virshup, 1999; Graves and Roach, 1995; Hoekstra et al., 1994). To test this prediction for Hhp1, we used synthetic, biotin-conjugated peptides corresponding to the unphosphorylated Hhp1 Cter and the fully phosphorylated Hhp1 Cter (Cter-6P). When linked to streptavidin beads, Hhp1 Cter-6P but not unphosphorylated Hhp1 Cter pulled down Hhp1ΔC from solution (Figure 2A). This result supports a direct, autophosphorylation-dependent interaction between the C-terminus and the kinase domain. We performed equilibrium binding assays at multiple concentrations of Cter to measure the dissociation constant of the Cter-kinase domain interaction. While the unphosphorylated Hhp1 Cter showed no appreciable binding even at the highest concentration, Hhp1 Cter-6P binding fit a one-site specific binding curve with a K_d_ = 240 μM (Figure 2B). In order to determine the binding affinity with both partners in solution, we performed isothermal titration calorimetry (ITC), and the K_d_ for Hhp1 Cter-6P determined via this method, 335 ± 74 μM, was consistent with the dissociation constant determined via pull-down (Figure 2C, Figure S3A). We detected no binding of the unphosphorylated Cter (Figure S3B).

**Figure 2:**
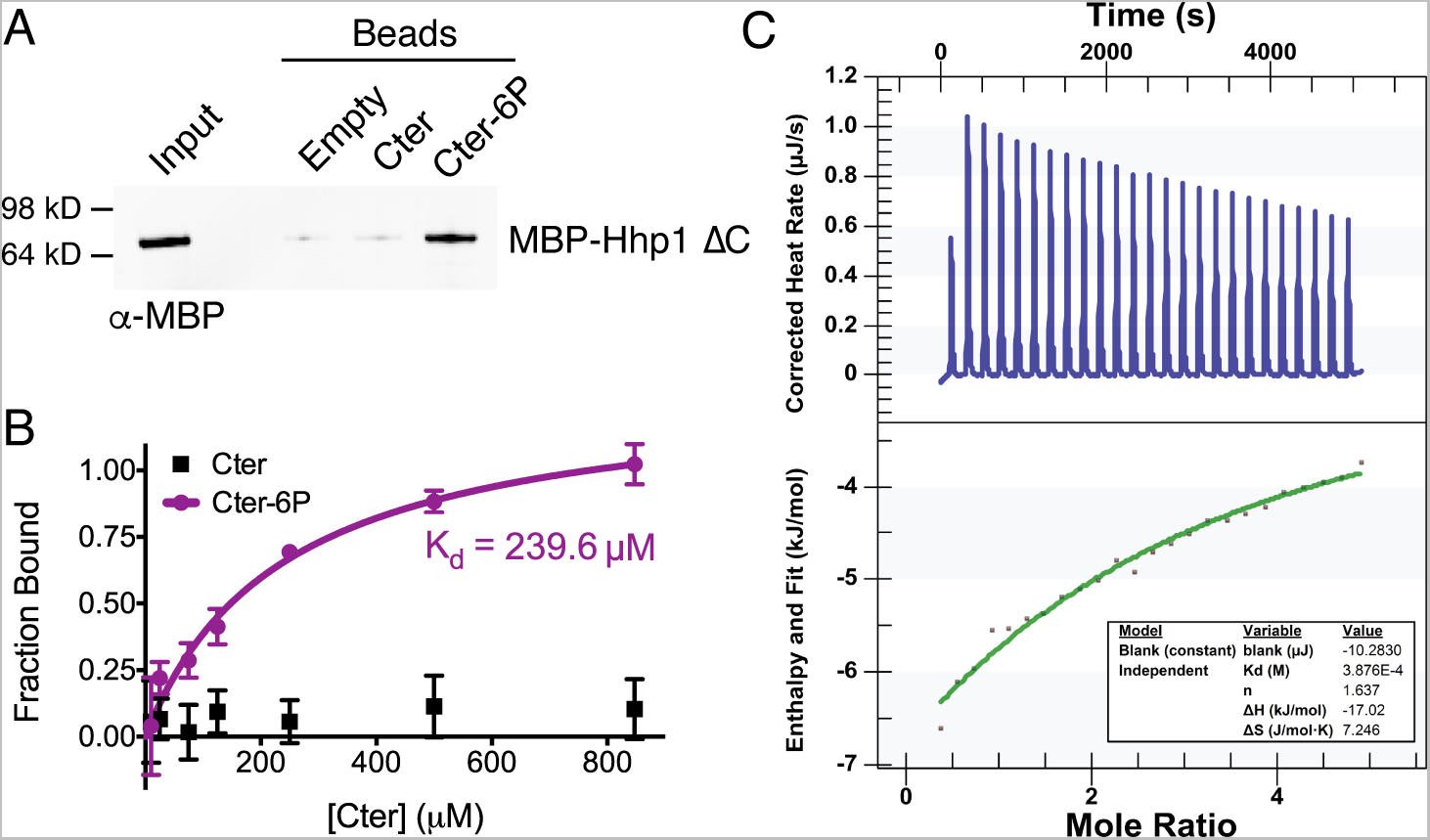
The Hhp1 C-terminus interacts with the kinase domain in a phosphorylation-dependent manner. (A) Hhp1 Cter or Cter-6P was conjugated to beads and incubated with MBP-Hhp1ΔC at 4°C for 1 h. Bound protein was visualized by western blot. Representative blot from three independent replicates. (B) Various concentrations of Hhp1 Cter or Cter-6P beads were incubated with MBP-Hhp1ΔC at 4°C for 1 h. Protein remaining in the supernatant was visualized by western blot and quantified by densitometry. Data from three independent replicates (mean ± SD) was fitted to a one-site specific binding curve to calculate the K_d_. (C) Representative ITC experiment with MBP-Hhp1ΔC and Cter-6P. Top panel shows raw data; bottom panel shows normalized integrated data. See Figure S3A-B for additional replicate and negative controls.

### The C-terminus of Hhp1 competes for binding to the substrate binding groove

To test whether the interaction between the phosphorylated Cter and the kinase domain inhibited substrate phosphorylation, we incubated Hhp1ΔC with increasing concentrations of the soluble Hhp1 Cter peptides and either the model substrate casein or the physiological substrate Rec11. Hhp1 Cter-6P inhibited substrate phosphorylation in a dose-dependent manner, with casein and Rec11 phosphorylation decreased by approximately half at the K_d_ of the binding interaction (Figure 3A-B). Similar to the binding data (Figure 2A-C), unphosphorylated Hhp1 Cter did not inhibit substrate phosphorylation.

**Figure 3:**
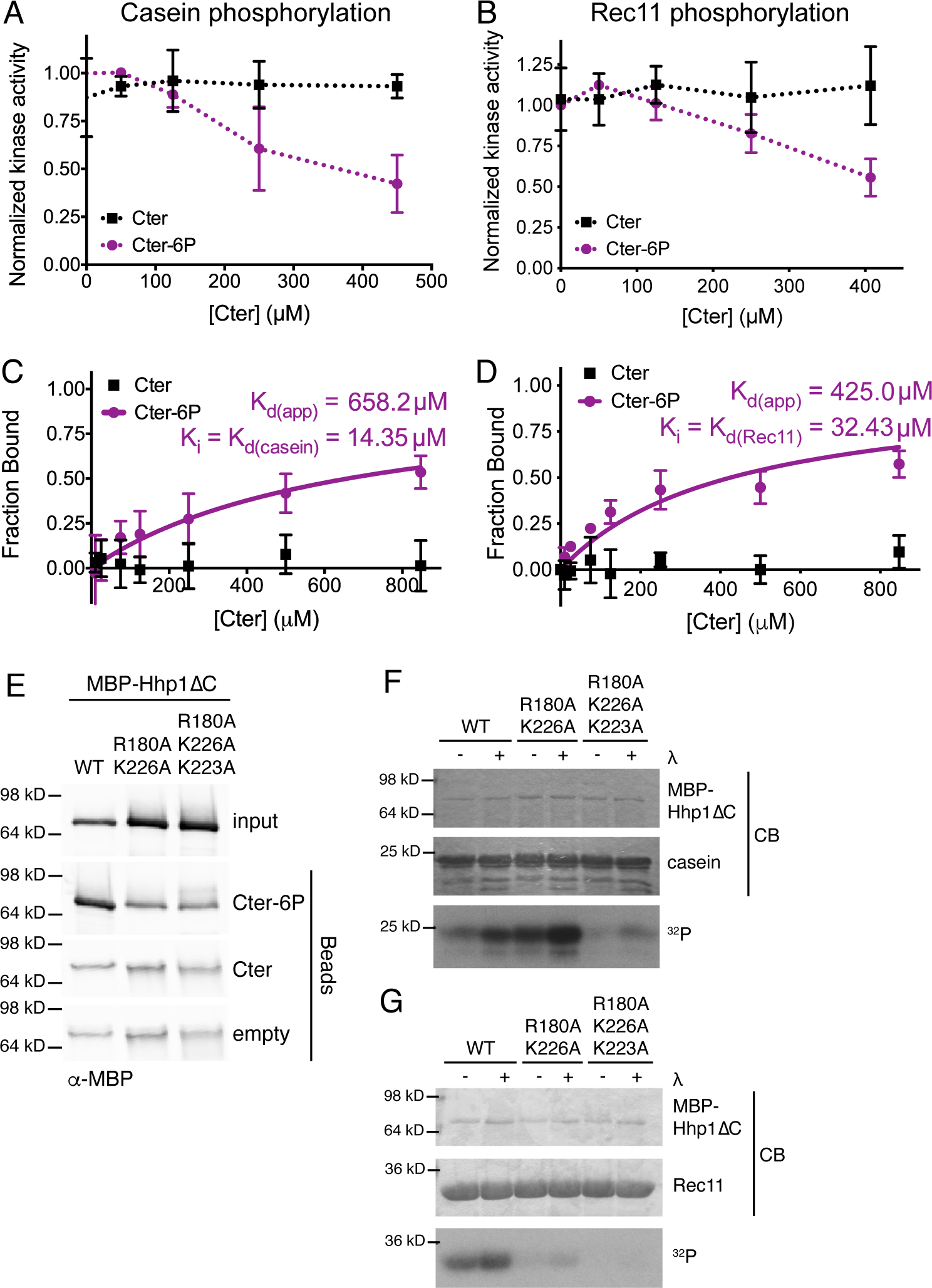
The Hhp1 C-terminus competes with substrates to bind the substrate binding groove. (A-B) Various concentrations of soluble Hhp1 Cter or Cter-6P were incubated with MBP-Hhp1ΔC, casein (A) or Rec11 (B), and γ-[^32^P]-ATP at 30°C for 1 h. Substrate phosphorylation was measured on a phosphorimager. Data from three independent replicates is shown as the mean ± SD. (C-D) Various concentrations of Hhp1 Cter or Cter-6P beads were incubated with MBP-Hhp1ΔC in the presence of 25 μM casein (C) or Rec11 (D) at 4°C for 1 h. Protein remaining in the supernatant was visualized by western blot and quantified by densitometry. Data from three independent replicates (mean ± SD) was fitted to a one-site specific binding curve to calculate the apparent K_d_ for the kinase domain-Cter interaction, and the substrate K_i_ was then calculated based on the change in K_d_ compared to that measured in Figure 2B. (E) Hhp1 Cter or Cter-6P was conjugated to beads and incubated with MBP-Hhp1ΔC with basic residues in the substrate binding groove mutated to alanine. Bound protein was visualized by western blot. Representative blot from three independent replicates. (F-G) MBP-Hhp1ΔC was treated +/- lambda phosphatase (λ), then incubated with casein (F) or Rec11 (G) and γ-[^32^P]-ATP at 30°C for 45 min. Substrate phosphorylation was detected by autoradiography (^32^P) and total protein by Coomassie (CB). Representative data from three independent replicates.

The pseudosubstrate model and structural studies predict that binding of the C-terminus to the CK1 kinase domain prevents substrate phosphorylation by directly competing with substrate for binding to the kinase domain (Gebel et al., 2020; Philpott et al., 2023). To confirm that this is the case in Hhp1, we repeated the equilibrium binding experiment in the presence of substrate (Figure 3C-D). Here, casein and Rec11 act as inhibitors to Cter binding, and we expected the apparent K_d_ for the Cter to increase due to competitive inhibition. Indeed, we found that the K_d(app)_ for Cter-6P in the presence of 25 μM casein was 658 μM, and the K_i(casein)_, which is equal to the K_d(casein)_, was 14 μM (Figure 3C). Similarly, the K_i(Rec11)_ = K_d(Rec11)_ = 32 μM (Figure 3D). It is noteworthy that the affinity of Hhp1ΔC for substrate is an order of magnitude greater than for the autophosphorylated C-terminus.

Because the substrate binding groove is positively charged, mutating basic residues in this region (Arg180, Lys223, and Lys226 in Hhp1) to alanine should prevent both Cter-6P binding and substrate phosphorylation. To test this, we expressed and purified recombinant Hhp1ΔC-R180A, K223A, K226A. To confirm that the overall secondary structure of the mutant kinase remained intact, we performed circular dichroism at far-UV wavelengths. The spectra of Hhp1ΔC-R180A, K223A, K226A resembled that of wildtype Hhp1ΔC, indicating that the mutations had not disrupted the catalytic domain structure (Figure S4A). However, these mutations eliminated binding to the Cter-6P peptide compared to wildtype Hhp1ΔC (Figure 3E) and dramatically reduced phosphorylation of casein (Figure 3F) and Rec11 (Figure 3G). These data are consistent with the proposal that the autophosphorylated C-termini of CK1s interact with the substrate binding grooves of the catalytic domains.

### Autophosphorylation-dependent, substrate-competitive tail binding is conserved in human CK1ε

To ask whether the autoinhibition model confirmed for Hhp1 was conserved in human orthologs, we first needed to identify the C-terminal autophosphorylation sites for a human CK1 enzyme. A previous study characterized CK1ε-MM2, which consisted of 8 proposed autophosphorylation sites mutated to alanine (Ser323, Thr325, Thr334, Thr337, Ser368, Ser405, Thr407, and Ser408; Gietzen and Virshup, 1999). We verified that these sites were autophosphorylated in vitro using phosphopeptide mapping; however, there were additional sites because the CK1ε-MM2 map was not identical to CK1εΔC (Figure S5A). Furthermore, CK1ε was found to incorporate up to 12 moles of phosphate per mole of enzyme (Cegielska et al., 1998), supporting the existence of additional phosphorylation sites. We immunoprecipitated CK1ε from HEK293 cells and identified additional candidate phosphorylation sites by mass spectrometry. As for Hhp1, the tail of CK1ε produces long phosphopeptides containing multiple serines and threonines, making phosphorylation site localization challenging. We expressed alanine substitution mutants at each candidate site and performed phosphopeptide mapping of the autophosphorylated, recombinant proteins to verify that sites were the result of autophosphorylation and to accurately localize sites (Figure S5A). Ultimately, we mutated an additional 7 sites (Ser343, Ser350, Thr351, Ser354, Ser389, Ser390, Ser391) in combination with the 8 sites in CK1ε-MM2 to generate CK1ε-15A (Figure 4A), which abolished all C-terminal autophosphorylation (Figure S5A). Similar to Hhp1, CK1εΔC can phosphorylate its C-terminus *in trans*, but not when these 15 sites are mutated to alanine (Figure 4B).

**Figure 4:**
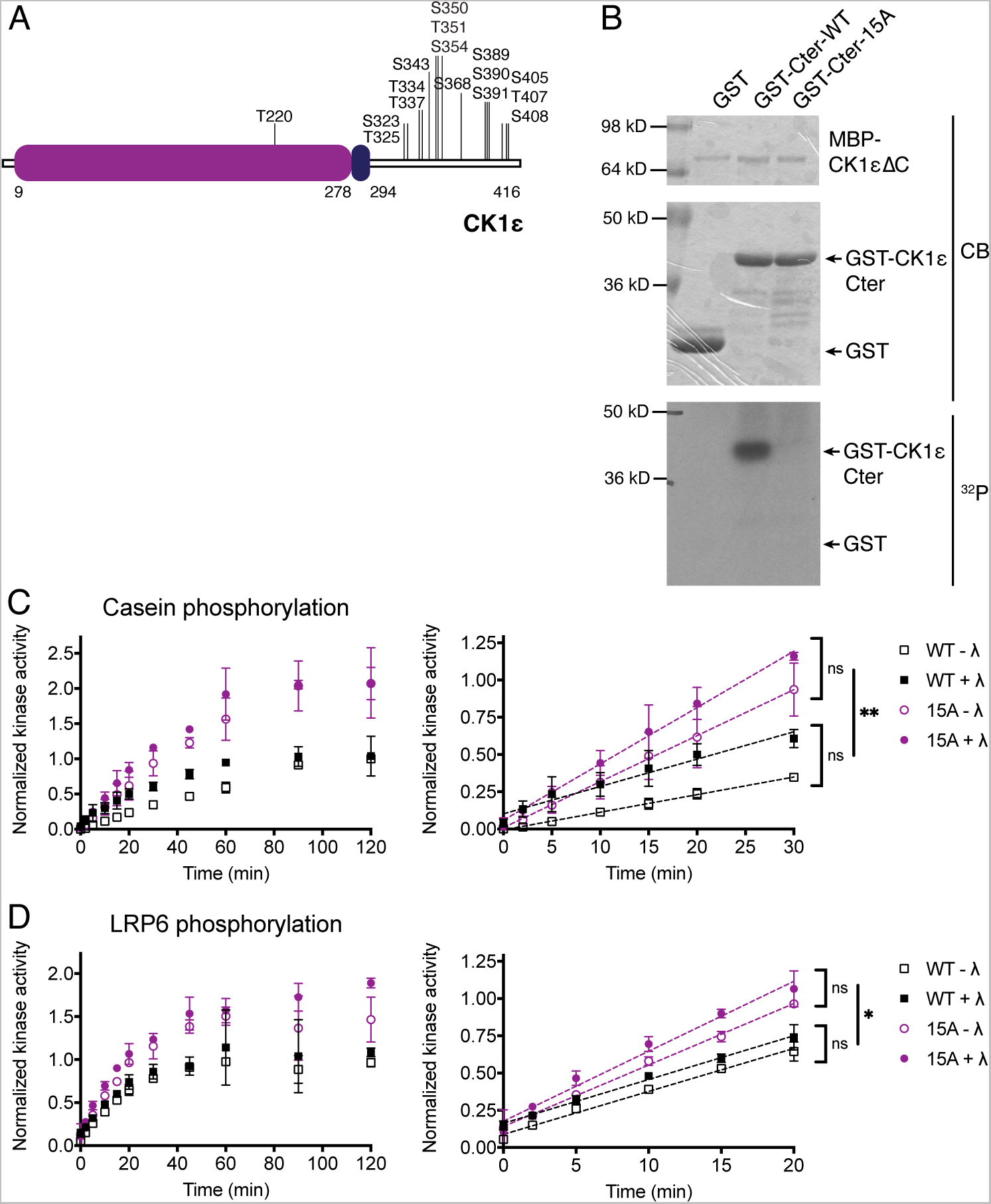
Autophosphorylation at 15 sites inhibits CK1ε activity. (A) Domain structure of human CK1ε showing confirmed autophosphorylation sites. (B) MBP-CK1εΔC was incubated with GST-CK1ε-Cter and γ-[^32^P]-ATP at 30°C for 2 h. Phosphorylation was detected by autoradiography (^32^P) and total protein by Coomassie (CB). (C-D) MBP-CK1ε-WT or -15A was treated +/- lambda phosphatase (λ), then incubated with substrate and γ-[^32^P]-ATP at 30°C. Reactions were quenched at timepoints from 0-120 min, and casein phosphorylation (C) or LRP6 phosphorylation (D) was measured on a phosphorimager. Data from three independent replicates is shown as the mean ± SD. The full timecourse is shown on the left; the initial rate was determined by the slope of the linear section of the curve shown on the right. ** = p < 0.001, * = p < 0.01, ns = not significant by one-way ANOVA of slopes.

CK1ε-15A phosphorylated two substrates, casein (Figure 4C) and the cytoplasmic domain of LRP6 (Figure 4D), a CK1ε substrate involved in Wnt signaling (Su et al., 2018; Swiatek et al., 2006; Wu et al., 2009; Zeng et al., 2005), at a greater rate than wildtype CK1ε, suggesting that autophosphorylation of the C-terminus at these 15 sites inhibits substrate phosphorylation.

We next tested the binding of CK1εΔC to the autophosphorylated C-terminus. Due to the length (122 amino acids) and number of phosphorylation sites, it was not possible to synthesize the entire autophosphorylated CK1ε Cter as we had for Hhp1 Cter. Instead, we used a peptide corresponding to the final 34 amino acids of CK1ε (called EC) due to the demonstrated effect of the extreme C-terminus on the phosphorylation of PER2 (Fustin et al., 2018; Narasimamurthy et al., 2018). The EC-6P peptide included the 6 autophosphorylation sites that occur in that region. Recombinant CK1εΔC interacted with EC-6P but not EC in a pulldown experiment (Figure 5A), and using ITC we determined that the K_d_ of this interaction was 343 ± 30 μM (Figure 5B, Figure S3C-D). Increasing concentrations of EC-6P inhibited LRP6 phosphorylation by CK1εΔC (Figure 5C).

**Figure 5:**
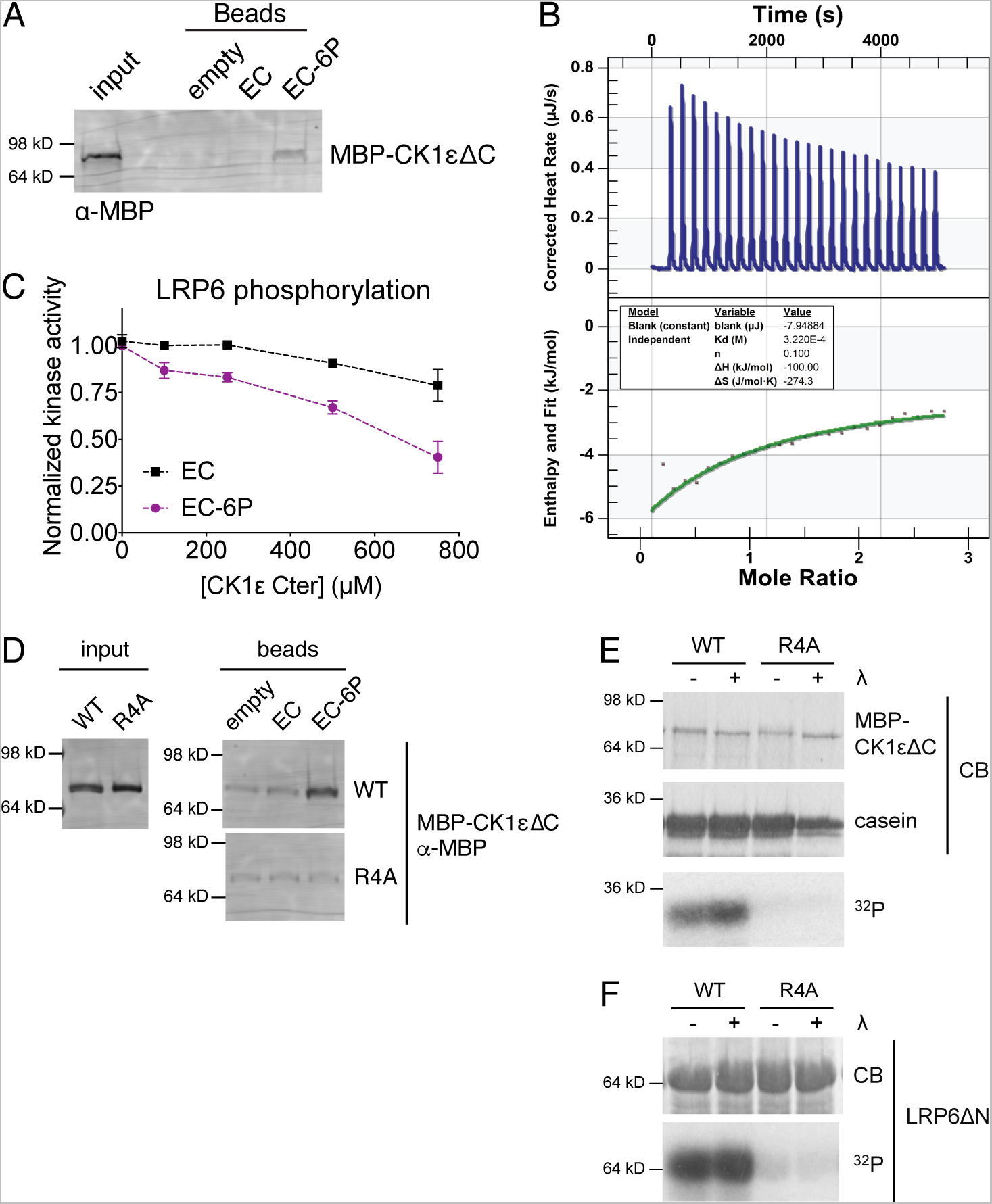
Phosphorylation-dependent tail binding and competition with substrate is conserved in human CK1ε. (A) CK1ε EC or EC-6P was conjugated to beads and incubated with MBP-CK1εΔC at 4°C for 1 h. Bound protein was visualized by western blot. Representative blot from two independent replicates. (B) Representative ITC experiment with MBP-CK1εΔC and EC-6P. Top panel shows raw data; bottom panel shows normalized integrated data. See Figure S3C-D for additional replicate and negative controls. (C) Various concentrations of soluble CK1ε EC or EC-6P were incubated with MBP-CK1εΔC, LRP6, and γ-[^32^P]-ATP at 30°C for 1 h. LRP6 phosphorylation was measured on a phosphorimager. Data from two independent replicates is shown as the mean ± SD. (D) CK1ε EC or EC-6P was conjugated to beads and incubated with MBP-CK1εΔC with basic residues in the substrate binding groove mutated to alanine (R4A). Bound protein was visualized by western blot. Representative blot from two independent replicates. (E-F) MBP-CK1εΔC was treated +/- lambda phosphatase (λ), then incubated with casein (E) or LRP6 (F) and γ-[^32^P]-ATP at 30°C for 45 min. Substrate phosphorylation was detected by autoradiography (^32^P) and total protein by Coomassie (CB). Representative data from two independent replicates.

Mutating basic residues in the substrate binding groove generated CK1εΔC-R178A, K221A, R222A, K224A, which was unable to interact with EC-6P (Figure 5D). This mutant also had dramatically reduced activity towards casein (Figure 5E) and LRP6 (Figure 5F) compared to wildtype, even though the secondary structure of the mutant was the same as wildtype (Figure S4B). We conclude that as for Hhp1, the autophosphorylated C-terminus of CK1ε competes with substrates to bind the catalytic domain.

### CK1 C-termini contribute to substrate specificity

The competitive binding interactions between the C-termini of Hhp1 and CK1ε and their substrates suggested that the C-termini might influence which substrates were targeted by each kinase. In other words, are all substrates equally competed by the phosphorylated C-termini, or are some more easily prevented from accessing the substrate binding groove than others? To answer this question, we measured the catalytic efficiency or specificity constant (k_cat_/K_m_) for the full-length and ΔC versions of Hhp1, Hhp2, CK1δ, and CK1ε on a few representative substrates (Figure 6A-D). In addition, we treated with lambda phosphatase to remove T220 autophosphorylation, which influences substrate specificity by changing the conformation of the substrate binding groove (Cullati et al., 2022). This allowed us to build a “specificity footprint” for each phospho-form of each kinase (Figure 6A-D).

**Figure 6:**
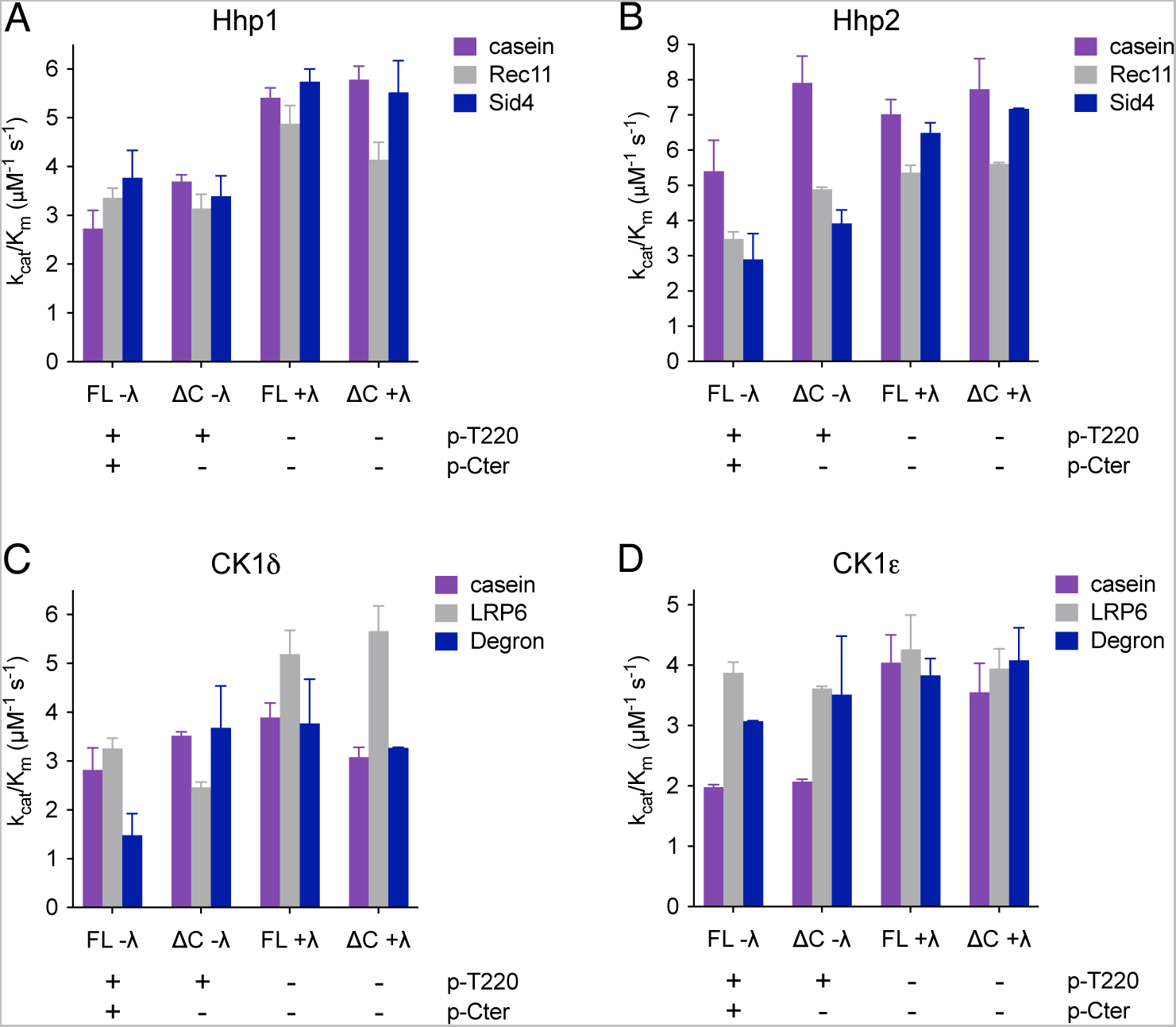
CK1 C-termini contribute to substrate specificity. Specificity footprints for Hhp1 (A), Hhp2 (B), CK1δ (C), and CK1ε (D). The indicated CK1 enzymes were treated +/- lambda phosphatase (λ), then incubated with the indicated substrates and γ-[^32^P]-ATP at 30°C. Reactions were quenched at timepoints from 0-60 min, and substrate phosphorylation was measured on a phosphorimager. The initial rate was determined by the slope of the linear section of the curve, then used to calculate the k_cat_/K_m_ (see Methods for details). Data from three independent replicates is shown as the mean ± SD.

To reduce complexity, the substrates that we chose did not require priming by other kinases. In addition to casein and Rec11, we used Sid4 for Hhp1 and Hhp2, which is a substrate involved in a yeast mitotic checkpoint (Johnson et al., 2013). For CK1δ and CK1ε, we used casein, LRP6, and a peptide from PER2 that includes the β-TrCP phosphodegron site (Philpott et al., 2023). We also tested whether truncating the C-termini affected the catalytic efficiency in the same way as abolishing the C-terminal autophosphorylation sites. We measured similar k_cat_/K_m_ values for Hhp1-6A and Hhp1ΔC (Figure S6A and Figure 6A) as well as for CK1ε-15A and CK1εΔC (Figure S6B and Figure 6D). This allowed us to compare the k_cat_/K_m_ for the different phospho-forms of Hhp2 and CKδ, even without knowing their C-terminal autophosphorylation sites.

We found that the catalytic efficiencies were different for different substrates – for example, full-length autophosphorylated Hhp1 was more likely to phosphorylate Sid4 than casein (Figure 6A). Removing either C-terminal autophosphorylation or T220 autophosphorylation tended to increase the k_cat_/K_m_ for all substrates, but the magnitude of this effect differed. For example, Hhp1ΔC had a ∼30% greater k_cat_/K_m_ for casein than Hhp1, but the efficiency of Rec11 and Sid4 phosphorylation was nearly the same for Hhp1ΔC and Hhp1, such that Hhp1ΔC had little preference for one substrate over the other (Figure 6A). Dephosphorylating the T220 sites further increased the k_cat_/K_m_ for casein and Sid4 but had a smaller effect on Rec11 (Figure 6A). This demonstrates that CK1 autophosphorylation state can influence not only the absolute values for k_cat_/K_m_, but importantly, also the relative values of substrates compared to each other.

Furthermore, different kinases had different catalytic efficiencies for the same substrate. Comparing Hhp2 to Hhp1, full-length autophosphorylated Hhp2 had a preference for casein over Rec11 and Sid4 (Figure 6B). Interestingly, removing Hhp2 autophosphorylation generated a pattern that resembled that of Hhp1 (Figure 6A-B), indicating that the two kinase isoforms have different substrate specificities when they are autophosphorylated, and these differences are diminished in the dephosphorylated forms. Similarly, for the human enzymes, the catalytic efficiencies differed between substrates, kinases, and phosphostates (Figure 6C-D). Consistent with previous data, removal of the CK1δv1 C-terminus greatly increased Degron phosphorylation (Isojima et al., 2009; Philpott et al., 2020), and the effect was more muted in CK1ε. Fully phosphorylated CK1δv1 had greater preference for casein and LRP6, while CK1ε had greater preference for LRP6 and the Degron. Dephosphorylation of both enzymes tended to increase catalytic efficiencies, and the dephosphorylated forms were more similar to each other, though CK1δ retained a preference for LRP6 (Figure 6C-D). These data suggest that across yeast and human enzymes, the effects of CK1 autophosphorylation depend on each kinase-substrate pair, and autophosphorylation of both T220 in the catalytic domain and the multiple sites in the C-termini of CK1s contribute to substrate specificity.

### Phosphorylation in the kinase domain influences C-terminal autophosphorylation rate and binding affinity

In light of these findings, we asked whether T220 autophosphorylation affected the probability of tail autophosphorylation and binding to the kinase domain. Preventing kinase domain autophosphorylation by mutating the T220 site increased the rate of C-terminal autophosphorylation in Hhp1-VV (Cullati et al., 2022) and CK1ε-T220N (Figure 7A) compared to the wildtype kinases. The localization of C-terminal autophosphorylation sites remained constant, however, because phosphopeptide maps of Hhp1-VV and CK1ε-T220N showed no change in C-terminal phosphopeptides (Figure S1C and Figure S5B).

**Figure 7:**
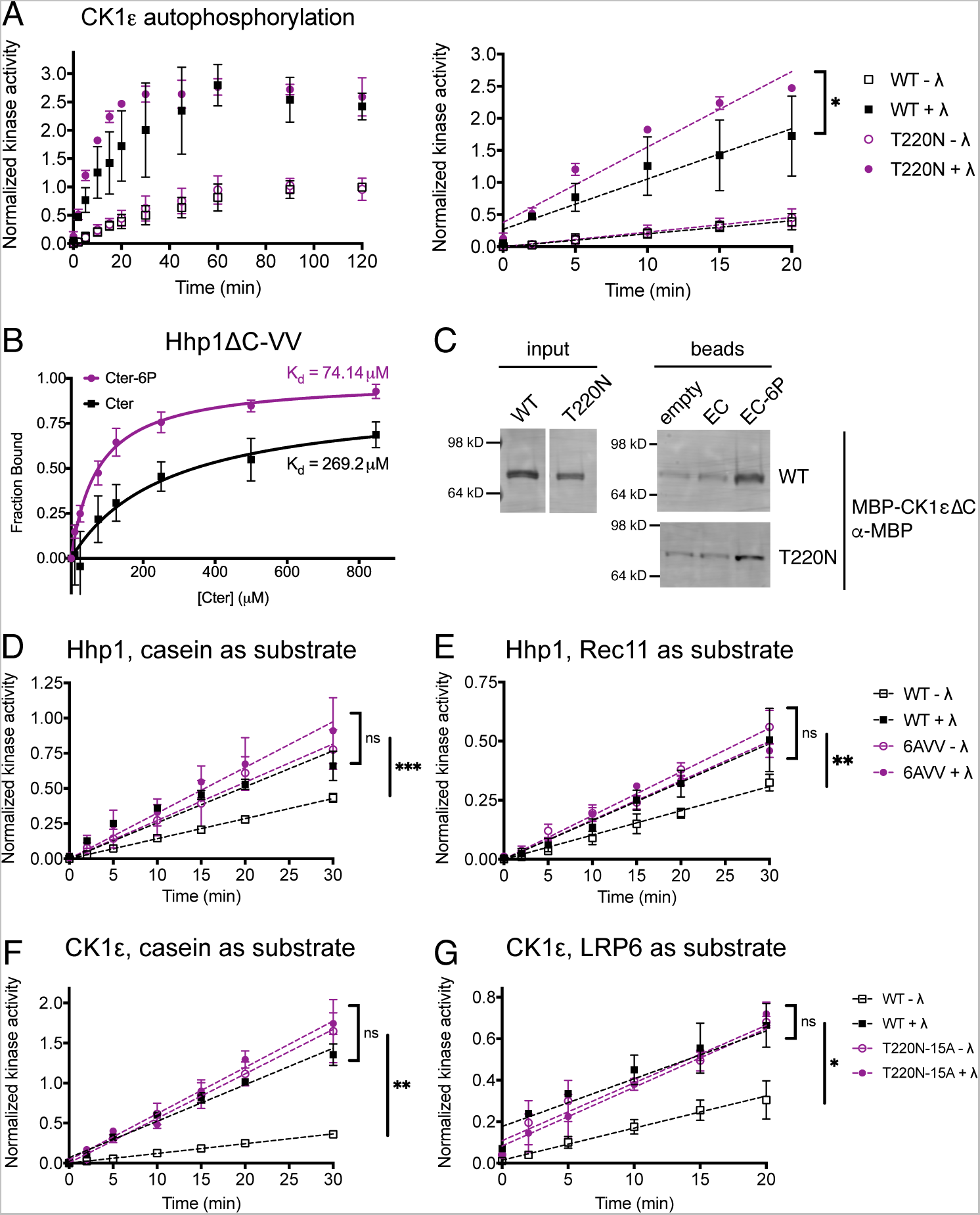
Phosphorylation in the kinase domain influences C-terminal autophosphorylation rate and binding affinity. (A) MBP-CK1ε was treated +/- lambda phosphatase (λ), then incubated with γ-[^32^P]-ATP at 30°C. Reactions were quenched at timepoints from 0-120 min, and autophosphorylation was measured on a phosphorimager. Data from three independent replicates is shown as the mean ± SD. The full timecourse is shown on the left; the initial rate was determined by the slope of the linear section of the curve shown on the right. * = p < 0.01 by one-way ANOVA of slopes. (B) Various concentrations of Hhp1 Cter or Cter-6P beads were incubated with MBP-Hhp1ΔC-VV at 4°C for 1 h. Protein remaining in the supernatant was visualized by western blot and quantified by densitometry. Data from three independent replicates (mean ± SD) was fitted to a one-site specific binding curve to calculate the K_d_. (C) CK1ε EC or EC-6P was conjugated to beads and incubated with MBP-CK1εΔC at 4°C for 1 h. Bound protein was visualized by western blot. Representative blot from two independent replicates. (D-E) MBP-Hhp1 was treated +/- λ, then incubated with substrate and γ-[^32^P]-ATP at 30°C. Reactions were quenched at timepoints from 0-120 min, and casein phosphorylation (D) or Rec11 phosphorylation (E) was measured on a phosphorimager. Data from three independent replicates is shown as the mean ± SD. *** = p < 0.0002, ** = p < 0.001, ns = not significant by one-way ANOVA of slopes. (F-G) MBP-CK1ε was treated +/- λ, then incubated with substrate and γ-[^32^P]-ATP at 30°C. Reactions were quenched at timepoints from 0-120 min, and casein phosphorylation (F) or LRP6 phosphorylation (G) was measured on a phosphorimager. Data from three independent replicates is shown as the mean ± SD. ** = p < 0.001, * = p < 0.01, ns = not significant by one-way ANOVA of slopes.

Hhp1ΔC-VV and CK1εΔC-T220N were still able to bind peptides corresponding to their phosphorylated C-termini (Figure 7B-C). The K_d_ for Hhp1ΔC-VV interacting with Cter-6P was 74 μM, approximately three-fold lower than for Hhp1ΔC (Figure 2B), and for the first time, we were able to detect binding of Hhp1ΔC-VV to the unphosphorylated Cter (Figure 7B). This suggests that catalytic domain phosphorylation decreases the likelihood that the Cter will be phosphorylated and therefore also the likelihood that the Cter will interact with the kinase domain. Further, catalytic domain phosphorylation increases the selectivity for binding phosphorylated Cter by preventing promiscuous interaction of the unphosphorylated tail. When both mechanisms were simultaneously disrupted in Hhp1-VV-6A and CK1ε-T220N-15A, these mutant kinases phosphorylated their substrates at a greater rate than wildtype, but the effects did not appear to be additive (Figure 7D-G), further supporting the idea that autophosphorylation in the C-terminus and autophosphorylation in the kinase domain are not independent of each other.

## Discussion

Although CK1 enzymes are multifunctional kinases involved in myriad essential signaling pathways related to human disease, the mechanisms by which their catalytic activities are regulated are not fully understood. Previous research has focused on extrinsic regulation of CK1 substrates by phosphatases and priming kinases (Bhandari et al., 2013; Ishiguro et al., 2010; Zeng et al., 2005). More recent work has also shown that CK1 activity can be modulated by binding interactions (Cruciat et al., 2013; Elmore et al., 2018), subcellular targeting (Elmore et al., 2018; Greer and Rubin, 2011), and a conserved autophosphorylation site in the kinase domain (Cullati et al., 2022). C-terminal autophosphorylation was known to inhibit CK1 activity (Cegielska et al., 1998; Gietzen and Virshup, 1999; Graves and Roach, 1995; Hoekstra et al., 1994), and it was proposed that phosphatases are required to activate CK1 in cells (Rivers et al., 1998). However, the biochemical model for how autophosphorylation inhibits CK1 had not been tested until recently.

We found that autophosphorylated C-termini can directly bind CK1 kinase domains to inhibit substrate phosphorylation. Our results support the idea that the C-terminus acts as a pseudosubstrate that can occupy the substrate binding groove in a manner similar to substrate peptides (Cegielska et al., 1998; Gebel et al., 2020; Gietzen and Virshup, 1999; Graves and Roach, 1995; Hoekstra et al., 1994; Philpott et al., 2023, 2020). The high substrate affinities that we measured are consistent with the product inhibition observed for the PER2 FASPS region and for p63 (Gebel et al., 2020; Philpott et al., 2023; Shinohara et al., 2017). These studies revealed that CK1 phosphorylates its substrates in a distributive manner, raising the questions of whether the C-termini are also phosphorylated in a distributive manner, if all sites are phosphorylated in a single molecule, and if there is any hierarchical phosphorylation of the C-termini, in which certain sites are preferentially phosphorylated.

The Cter interaction with the kinase domain has a relatively high K_d_ (Figure 2B-C, Figure 5B). In vitro, the binding affinity was necessarily measured *in trans*, but in vivo, the C-terminus is attached to the kinase domain. Previous work showed that autophosphorylation of Hhp1 (Hoekstra et al., 1994), rat CK1δ (Graves and Roach, 1995), human CK1α (Cegielska and Virshup, 1993) and human CK1ε (Cegielska et al., 1998) occurs *in cis.* In this configuration, the Cter would be at a very high local concentration, which would favor its phosphorylation and overcome the low affinity observed in vitro. Another contributing factor could be that the autophosphorylation sites are not located in sequences that conform to the canonical CK1 motif (Flotow et al., 1990; Flotow and Roach, 1991; Johnson et al., 2023). Other known substrates that harbor non-consensus CK1 sites are phosphorylated at slower rates than consensus sites, for example the FASP site on PER2 that is phosphorylated by CK1δv2 and CK1ε (Narasimamurthy et al., 2018; Philpott et al., 2020).

We found that substrates have much higher affinities for the kinase domain than the phosphorylated C-terminus. Indeed, substrates can competitively inhibit Cter binding even when all autophosphorylation sites are occupied in a synthesized phosphopeptide present at very high concentration (Figure 3C-D). This would mean that CK1s are never fully inhibited by this mechanism in cells, which makes biological sense if they are being used in multiple processes at multiple times. We propose a displacement specificity model (Figure 8) in which truncation or dephosphorylation of the tail would not be strictly required for all substrate phosphorylation events. Substrates that have a high affinity or that are present at a high concentration may instead out-compete the tail for access to the kinase active site. The action of phosphatases on the tail may be utilized as an additional level of regulation to allow scarce or low-affinity substrates to be phosphorylated under specific conditions. There is evidence that PP1 (Cullati et al., 2022; Rivers et al., 1998), PP2A (Rivers et al., 1998), calcineurin (Liu et al., 2002), and PP5 (Partch et al., 2006) can dephosphorylate CK1ε to increase its activity. Future in vivo studies could address which substrates and signaling pathways require phosphatase(s) to activate CK1 and which may be independent. Furthermore, CK1 C-termini differ in length and sequence, yet they all appear to interact with the conserved kinase domain; this would imply that different tail isoforms have different affinities for the kinase domain, allowing different substrates to displace each one.

**Figure 8:**
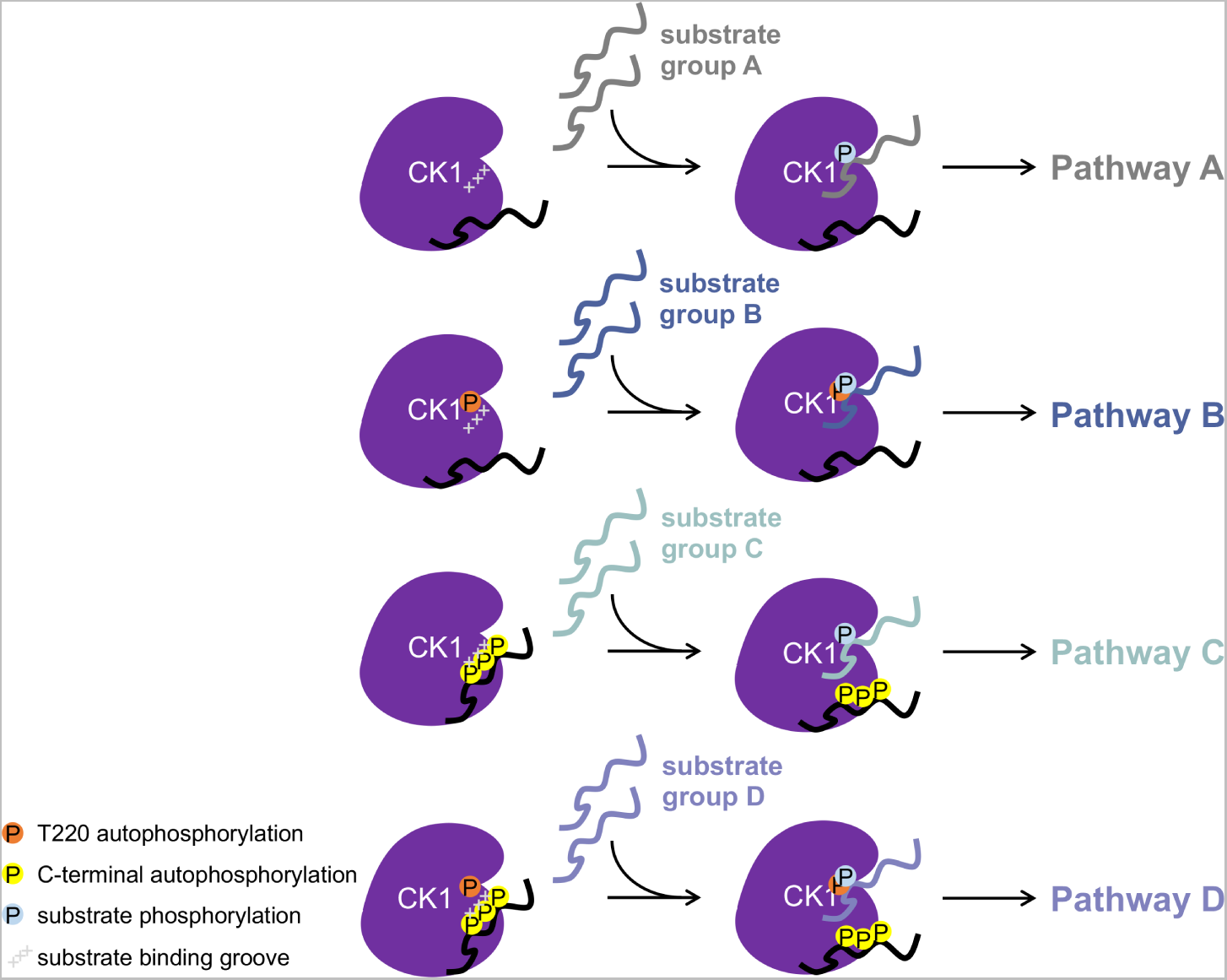
Displacement specificity model for the regulation of CK1 substrate specificity by autophosphorylation. At least four different phospho-forms of CK1 may exist. Depending on the conformation of the substrate binding groove (mediated by kinase domain autophosphorylation, orange) and the affinity of the C-terminus for the kinase domain (mediated by C-terminal autophosphorylation, yellow), different groups of substrates may be acted upon by each phospho-form (substrate phosphorylation, blue). If these substrates are involved in different signaling pathways, autophosphorylation could serve to stratify these substrates and direct CK1 activity to the proper role at the proper time.

The displacement specificity model also accounts for conformational changes in the substrate binding groove induced by autophosphorylation in the catalytic domain (Cullati et al., 2022). We suggest that there are at least four different phospho-forms of CK1 based on occupancy of catalytic domain and tail autophosphorylation sites, each of which would have a pool of preferred high-affinity substrates (Figure 8). If these substrate pools are involved in different signaling pathways, this may be a mechanism to stratify substrates. Within cells, phosphatases and other kinases also target CK1 and affect its function (Bischof et al., 2013; Giamas et al., 2007; Ianes et al., 2016; Meng et al., 2016). It will be informative to understand how CK1 is phosphorylated and dephosphorylated in different cellular contexts. The displacement specificity model provides a framework for these studies by integrating biochemical mechanism with cellular signaling outputs, and it can be continually tested with new substrates as we uncover more about CK1 signaling.

## Materials and Methods

### Molecular biology and protein purification

All plasmids were generated by standard molecular biology techniques. For protein production, cDNA for Hhp1, Hhp2, CK1δv1, and CK1ε was cloned into the pMAL-C2 vector. CK1ΔC constructs consisted of the following amino acids: Hhp1 1-296, Hhp2 1-295, CK1δ 1-294, CK1ε 1-294. Mutagenesis was performed using QuikChange XL and QuikChange Multi site-directed mutagenesis kits (Agilent Technologies). Plasmids were validated by DNA sequencing.

Protein production was induced in *Escherichia coli* Rosetta2(DE3)pLysS cells by addition of 0.1 mM IPTG overnight at 17°C. Cells were lysed using 300 µg/mL lysozyme for 20 min followed by sonication. MBP fusion proteins were purified on amylose beads (New England Biolabs) in column buffer (20 mM Tris pH 7.4, 150 mM NaCl, 1 mM EDTA, 0.1% NP-40, 1 mM DTT, 1 mM PMSF, 1.3 mM benzamidine, protease inhibitor tablets [Roche]) and eluted with maltose (20 mM Tris pH 7.4, 150 mM NaCl, 1 mM EDTA, 1 mM DTT, 1 mM PMSF, 1.3 mM benzamidine, 10 mM maltose, 10% glycerol). For purification of Sid4, Sid4 cDNA was cloned into the pET His6 MBP PreScission LIC cloning vector (gift from Scott Gradia, Addgene plasmid #29721). His-MBP-Sid4 was purified on amylose beads in column buffer containing 300 mM NaCl, 20 mM MgCl_2_, and 20 mM Na_2_SO_4_. On-bead cleavage with PreScission (GE Healthcare) protease was performed in cleavage buffer (50 mM Tris pH 7.0, 300 mM NaCl, 1 mM EDTA, 0.1% NP-40, 1 mM DTT, 10% glycerol, 20 mM MgCl_2_, 20 mM Na_2_SO_4_) for 16 h at 4°C using 1 U protease per 200 µg fusion protein. Protease was removed by incubation with GST-Bind beads (EMD Millipore). Sid4 was concentrated using Amicon Ultra centrifugal filters (Millipore). GST fusion proteins were purified on GST-Bind beads in GST Bind buffer (10 mM NaPO_4_ pH 7.3, 150 mM NaCl, 2.7 mM KCl, 0.1% NP40, 1 mM EDTA, 1mM DTT, 1 mM PMSF, 1.3 mM benzamidine, protease inhibitor tablets [Roche]) and eluted with glutathione (50 mM Tris pH 8.0, 100 mM NaCl, 1 mM DTT, 1 mM PMSF, 1.3 mM benzamidine, 10 mM glutathione, 10% glycerol). The human PER2 Degron sequence (amino acids 475-505; Philpott et al., 2023) was synthesized as a gBlock (IDT), cloned into pMAL-C2, and expressed and purified as above. GST-Rec11-N3 and MBP-LRP6ΔN were expressed and purified as previously described (Cullati et al., 2022). Casein was dephosphorylated alpha casein purchased from Sigma.

### In vitro kinase assays

Kinases were treated with 0.5 µL lambda phosphatase (New England Biolabs) per 1 µg protein for 45 min at 30°C in PMP buffer (New England Biolabs) plus 1 mM MnCl_2_. For negative controls, an equivalent volume of buffer was added instead of phosphatase. Phosphatase reactions were quenched by the addition of 8 mM Na_3_VO_4_ immediately prior to the kinase assay. All kinase assays were performed at 30°C and quenched by boiling in SDS-PAGE sample buffer. Proteins were separated by SDS-PAGE and stained in Coomassie to visualize total protein, then gels were dried prior to detection of phosphoproteins.

Phosphorylation of the Hhp1 and CK1ε Cter was performed with 1 µg MBP-Hhp1ΔC or 1 µg MBP-CK1εΔC and 4 µg GST-Hhp1 Cter or GST-CK1ε Cter in PMP buffer plus 250 μM cold ATP, 1 µCi γ-[^32^P]-ATP, and 10 mM MgCl_2_ for 2 h. Phosphorylation of casein and Rec11 by Hhp1 and LRP6 by CK1ε mutants was performed with 300 ng kinase and 25 µM substrate in PMP buffer plus 100 μM cold ATP, 1 µCi γ-[^32^P]-ATP, and 10 mM MgCl_2_ for 45 min. Phosphorylated proteins were visualized by autoradiography.

Phosphorylation of casein and Rec11 in the presence of Hhp1 Cter and LRP6 in the presence of CK1ε Cter was performed with 200 ng MBP-Hhp1ΔC or MBP-CK1εΔC, 25 µM substrate, and 0-455 μM Cter peptide in PMP buffer plus 250 μM cold ATP, 1 µCi γ-[^32^P]-ATP, and 10 mM MgCl_2_ for 1 h. Kinetic assays were performed using 200 ng MBP-Hhp1 or MBP-CK1ε and 25 µM substrate in PMP buffer plus 250 μM cold ATP, 1 µCi γ-[^32^P]-ATP, and 10 mM MgCl_2_. Reactions were quenched at time points from 0-120 min. Phosphorylated proteins were visualized using an FLA7000IP Typhoon Storage Phosphorimager (GE Healthcare Life Sciences) and quantified in ImageJ. Relative kinase activity was plotted in Prism.

To determine the k_cat_/K_m_, 200 ng MBP-tagged kinase (0.12 µM for full-length and 0.13 µM for ΔC) and 0.62 µM substrate were incubated in PMP buffer plus 250 μM cold ATP, 2 µCi γ-[^32^P]-ATP, and 10 mM MgCl_2_. Reactions were quenched at time points from 0-60 min. Phosphorylated proteins were visualized using an FLA7000IP Typhoon Storage Phosphorimager, quantified in ImageJ, and scaled to the moles of substrate. At this substrate concentration, [S] << K_m_ and [E_]free_ approximately equals [E]_T_. Therefore, the measured reaction rate (V_o_) equals (k_cat_/K_m_)[E]_T_[S].

### In vitro binding assays

Peptides corresponding to amino acids 297-365 of Hhp1 were synthesized and linked to biotin at the N-terminus, either unphosphorylated (Cter) or phosphorylated at S326, S327, T345, T346, S354, and T356 (Cter-6P), by Bio-Synthesis, Inc. Peptides corresponding to amino acids 383-416 of CK1ε were synthesized and linked to biotin at the N-terminus, either unphosphorylated (EC) or phosphorylated at S389, S390, S391, S405, T407, and S408 (EC-6P), by LifeTein LLC. Peptides were conjugated to streptavidin sepharose (Amersham Biosciences) at a concentration of 6 μg peptide per 1 μL bead slurry by rotating at 4°C overnight in binding buffer (50 mM Tris pH 7.0, 25 mM NaCl, 1 mM EDTA, 1 mM DTT). Beads were washed to remove unbound peptide, and the concentration of bound peptide was confirmed by SDS-PAGE. In all experiments, unconjugated streptavidin sepharose was used as a negative control (“Empty”), and beads were blocked in 5% BSA by rotating at 4°C for 30 min prior to binding assays in order to reduce nonspecific binding.

Peptide pull-downs were performed by incubating 2 μg MBP-Hhp1ΔC or MBP-CK1εΔC with 500 μM peptide in binding buffer for 1 h at 4°C, rotating. Beads were washed to remove unbound kinase, then boiled in SDS-PAGE sample buffer to elute. Bound protein was detected by western blot using mouse anti-MBP diluted 1:10,000 (New England Biolabs), fluorescent secondary antibodies (Li-Cor Biosciences), and an Odyssey CLx (Li-Cor Biosciences).

Equilibrium binding assays were performed according to (Pollard, 2010): The volume of peptide-conjugated beads was adjusted to yield peptide concentrations of 0-847 μM in the final reaction volume of 50 μL. Unconjugated streptavidin sepharose was then added to equalize the bead volume of all reactions. Beads were incubated with 100 ng MBP-Hhp1ΔC or 50 ng MBP-CK1εΔC in binding buffer for 1 h at 4°C, rotating. The supernatants were removed, and unbound kinase was detected by western blot as above. The unbound fraction was quantified on the Odyssey CLx, background subtracted, and normalized to the 0 μM condition. The fraction bound (1 – unbound) was plotted in Prism and fit to a one-site specific binding equation to calculate the K_d_. Competitive binding experiments were performed as above, but 25 μM substrate was added to the beads along with the kinase. The one-site specific binding equation was constrained such that the Bmax = 1 to calculate the apparent K_d_. The K_i_ was determined by the equation below, where [I] = [substrate] = 25 μM:

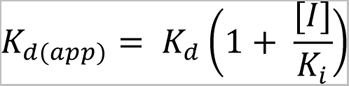

### Isothermal titration calorimetry

ITC measurements were conducted on a Nano-ITC instrument (TA Instruments) in ITC buffer (10 mM NaPO_4_ pH 7.5, 150 mM NaCl, 0.25 mM TCEP). The sample cell was filled with 300 μL of 100 μM MBP-Hhp1ΔC or MBP-CK1εΔC, while the syringe contained 50 μL of 2 mM peptide (Cter, Cter-6P, EC, or EC-6P). All solutions were degassed prior to being loaded into the cell. Aliquots (2 μL) of peptide solutions were injected into CK1 protein solutions at 25°C with an interval gap of 200 s and the syringe rotating at 150 rpm to ensure proper mixing. Data were analyzed using Nanoanalyser software to extract the thermodynamic parameters; the K_d_ was obtained after fitting the integrated and normalized data to a single-site binding model. Experiments were performed in duplicate to ensure reproducibility (see Figures 2C, 5B, and S3A-D).

### Yeast methods

*S. pombe* strains used in this study (Table S1) were grown in yeast extract (YE) media (Moreno et al., 1991). For *hhp1* gene replacements, haploid *hhp1::ura4+* was transformed using standard lithium acetate methods (Keeney and Boeke, 1994) to introduce linear *hhp1-6A* and *hhp1-6N* gene fragments generated by digestion of pIRT2-hhp1-6A and pIRT2-hhp1-6N plasmids with BamHI and PstI. Integrants were selected based on resistance to 1.5 mg/ml 5-fluoroorotic acid (Fisher Scientific) and validated by whole-cell PCR using primers homologous to endogenous flanking sequences in combination with those within the ORF. Truncation of *hhp1* was accomplished by insertion of the *kanMX6* cassette from pFA6 as previously described (Bähler et al., 1998), followed by selection on G418 (100 µg/mL; Sigma-Aldrich, St. Louis, MO). Tagged Hhp1-6N was generated by insertion of the *mNeonGreen:kanMX6* or *3xFLAG:kanMX6* cassettes from pFA6 as previously described (Bähler et al., 1998), followed by selection on G418. Insertions were validated by whole-cell PCR using primers homologous to the resistance cassette and the endogenous ORF. Diploid strains were constructed using standard techniques (Moreno et al., 1991); intragenic complementation between *ade6-M210* and *ade6-M216* alleles and maintenance on *ade^-^* growth medium ensured strains remained stable. All constructs and integrants were sequenced to verify their accuracy. *S. pombe* genomic sequences and annotation from PomBase (Harris et al., 2022).

For serial dilution growth assays, cells were cultured in YE at 32°C until mid-log phase, three 10-fold serial dilutions starting at 4x10^6^ cells/mL were made, 4 μL of each dilution was spotted on YE plates or YE plates containing 7.5 mM hydroxyurea (Sigma-Aldrich, St. Louis, MO), and cells were grown at the indicated temperatures for 3 d. For live cell imaging, cells were grown at 25°C. Images were acquired with a Personal DeltaVision microscope system (Leica Microsystems) that includes an Olympus IX71 microscope, 60x and 100x NA 1.42 PlanApo oil immersion objectives, a pco.edge sCMOS camera, and softWoRx imaging software. Images in figures are deconvolved maximum intensity projections of z sections spaced at 0.5 µm.

For immunoprecipitations, cell pellets were snap frozen then lysed by bead disruption using a FastPrep cell homogenizer (MP Biomedicals) in NP-40 buffer under denaturing conditions as previously described (Gould et al., 1991), except with the addition of protease inhibitor tablets (Roche) and phosphatase inhibitor tablets (Roche). Hhp1-3xFLAG was immunoprecipitated using 2 μg FLAG-M2 (Sigma). Proteins were separated by SDS-PAGE, transferred to Immobilon-P polyvinylidene fluoride membrane (Millipore), and immunoblotted with FLAG-M2 (1 μg/mL) followed by fluorescent anti-mouse secondary antibody (Li-Cor Biosciences) and imaging on an Odyssey CLx (Li-Cor Biosciences).

### Mass spectrometry

TCA-precipitated proteins were subjected to mass spectrometric analysis on an LTQ Velos (Thermo) by 3-phase multidimensional protein identification technology (MudPIT) as previously described (Chen et al., 2013) with the following modifications. Proteins were resuspended in 8 M urea buffer (8 M urea in 100 mM Tris pH 8.5), reduced with Tris (2-carboxyethyl) phosphine, alkylated with 2-chloro acetamide, and digested with trypsin or elastase. The resulting peptides were desalted by C-18 spin column (Pierce). For the kinase assay samples 6 salt elution steps were used (i.e. 25, 50, 100, 600, 1000, and 5000 mM ammonium acetate). Raw mass spectrometry data were filtered with Scansifter and searched by SEQUEST. Scaffold (version 3.6.0 or version 4.2.1) and Scaffold PTM (version 3.0.1) (both from Proteome Software, Portland, OR) were used for data assembly and filtering. The following filtering criteria were used: minimum of 90.0% peptide identification probability, minimum of 99% protein identification probability, and minimum of two unique peptides.

### Phosphopeptide mapping

Autophosphorylation reactions were performed with 4 µg lambda phosphatase-treated kinase in PMP buffer plus 100 μM cold ATP, 4 µCi γ-[^32^P]-ATP, and 10 mM MgCl_2_ at 30°C for 30 min. Reactions were quenched by boiling in SDS-PAGE sample buffer and proteins were separated by SDS-PAGE. Phosphorylated proteins were transferred to PVDF membranes. Proteins were digested off the membrane with 10 µg trypsin at 37°C overnight. Peptides were lyophilized and resuspended in pH 1.9 buffer. Tryptic peptides were separated in the first dimension by thin-layer electrophoresis and in the second dimension by chromatography (Boyle et al., 1991). After separations, TLC plates were exposed to film for 2-4 d at -80°C with intensifying screens.

### Circular dichroism

Recombinant MBP-Hhp1ΔC and MBP-CK1εΔC proteins were dialyzed into CD buffer (10 mM sodium phosphate pH 7.4, 150 mM Na_2_SO_4_, 0.5 mM EDTA, 0.5 mM DTT) overnight at 4°C. Proteins were diluted to 100 ng/µL in CD buffer and measured at RT in a 1 mm path length cuvette on a Jasco J-810 spectropolarimeter. Far-UV measurements were collected over a range of 260-190 nm at 0.5 nm resolution, and three scans were averaged for each spectrum. A blank consisting of CD buffer was subtracted from each protein spectrum, and the data (mdeg) were converted to mean residue ellipticity.

## Acknowledgments

We acknowledge technical assistance from Yufan Shan and Alaina Willet for the Hhp1-6N-mNG imaging in Fig. S2C. We thank members of the Gould laboratory for helpful discussions and critical comments on the manuscript, especially Rodrigo Guillen and Eric Zhang for their assistance with the CK1 project. **Funding**: S.N.C. was supported by the Integrated Biological Systems Training in Oncology Program (T32-CA119925). This work was funded by R35-GM131799 to K.L.G. **Author contributions**: S.N.C. and K.L.G. conceptualized the project and wrote the manuscript. Experiments were performed by J.S.C., K.A., and S.N.C. **Competing interests**: The authors declare no conflicts of interest. **Data and materials availability**: All data are available in the main text or the supplementary materials. All regents are freely available upon request.

## Supplementary Material

Figure S1: Identification of Hhp1 autophosphorylation sites.

Figure S2: Characterization of *hhp1* C-terminal autophosphorylation site mutants.

Figure S3: Hhp1 and CK1ε do not bind unphosphorylated tail peptides. Binding

Figure S4: Mutating the substrate binding groove does not disrupt kinase domain structure.

Figure S5: Identification of CK1ε autophosphorylation sites Figure S6: Specificity footprints of Hhp1-6A and CK1ε-15A.

Table S1: S. pombe strains used in this study.

## Notes

### Competing Interest Statement

The authors have declared no competing interest.

