## Supplemental Figures 1-6 and Supplemental Table 1 for "Substrate displacement of CK1 C-termini regulates kinase specificity"

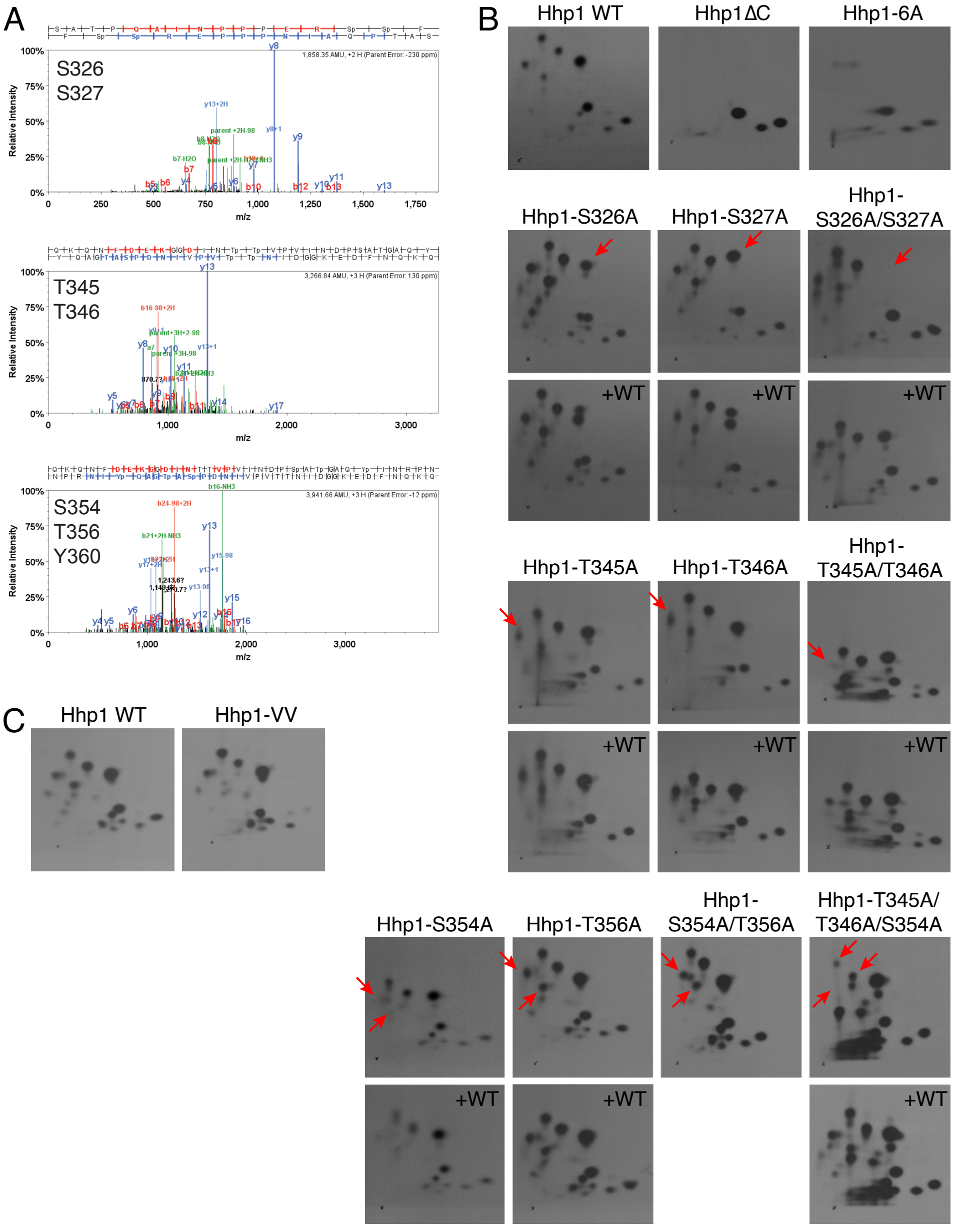


Supplemental Figure 1

**Figure S1: Identification of Hhp1 autophosphorylation sites.** (A) Recombinant MBP-Hhp1 was analyzed for post-translational modifications by mass spectrometry. Representative spectra of candidate autophosphorylation sites are shown. (B) The indicated Hhp1 mutants was treated with lambda phosphatase, then incubated with $\gamma$-[$^{32}$P]-ATP at 30°C for 30 min. Proteins were digested with trypsin, and peptides were separated by thin-layer electrophoresis and chromatography. Phosphopeptides were detected by autoradiography. Red arrows point out phosphopeptides affected by alanine substitutions. (C) Phosphopeptide map of C-terminal autophosphorylation sites in MBP-Hhp1-VV.

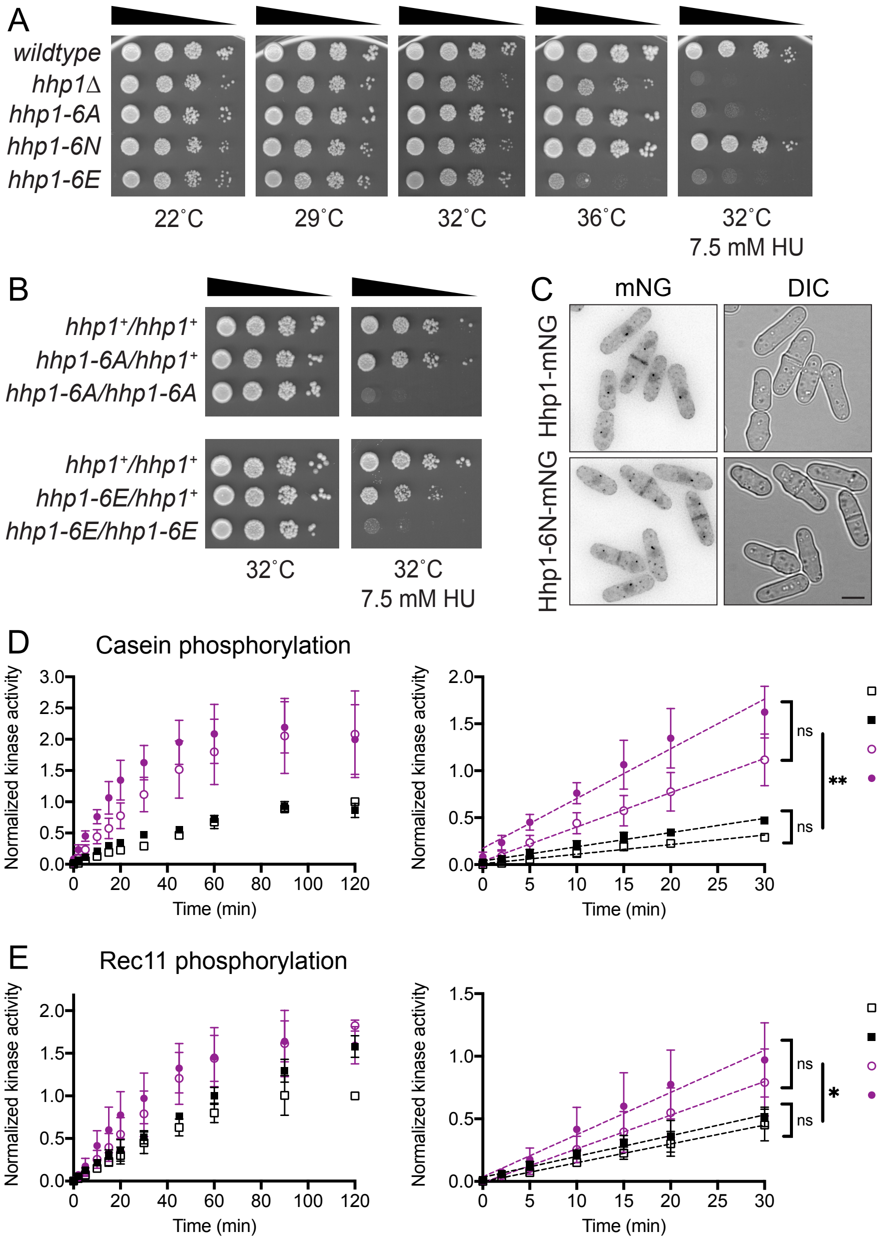


Supplemental Figure 2

**Figure S2: Characterization of *hhp1* C-terminal autophosphorylation site mutants.** (A) Mutant alleles were integrated at the endogenous *hhp1* locus, and 10-fold serial dilutions were spotted on YE with and without 7.5 mM hydroxyurea (HU). Cells were grown for 3 d at 32°C. Representative plates from 3 independent replicates are shown. (B) Diploid strains demonstrating *hhp1-6A* recessive loss-of-function. (C) Localization of fluorescently-tagged Hhp1-6N-mNG. (D-E) MBP-Hhp1-6N was treated +/- lambda phosphatase ($\lambda$), then incubated with substrate and $\gamma$-[$^{32}$P]-ATP at 30°C. Reactions were quenched at timepoints from 0-120 min, and casein phosphorylation (D) or Rec11 phosphorylation (E) was measured on a phosphorimager. Data from three independent replicates is shown as the mean ± SD. *** = $p < 0.0002$, ** = $p < 0.001$, ns = not significant by one-way ANOVA of slopes.

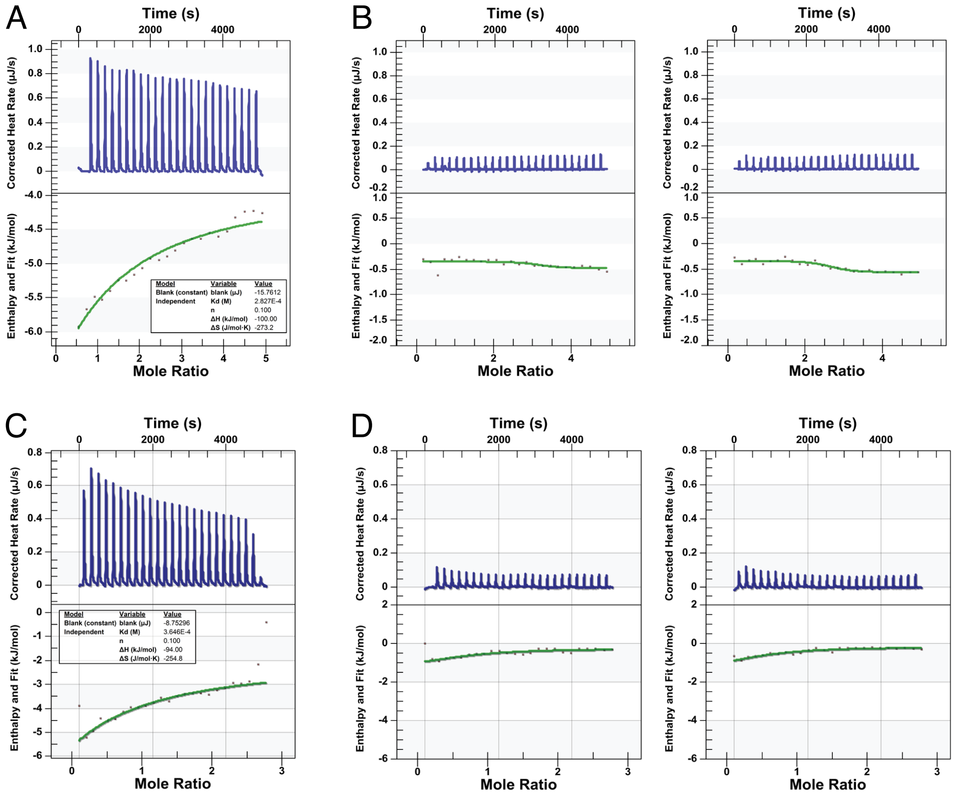


**Figure S3: Hhp1 and CK1ε do not bind unphosphorylated tail peptides.** Binding affinities determined by ITC. Top panel shows raw data; bottom panel shows normalized integrated data. (A) Second replicate of MBP-Hhp1ΔC binding Cter-6P. (B) MBP-Hhp1ΔC incubated with Cter. (C) Second replicate of MBP-CK1εΔC binding EC-6P. (B) MBP-CK1εΔC incubated with EC.

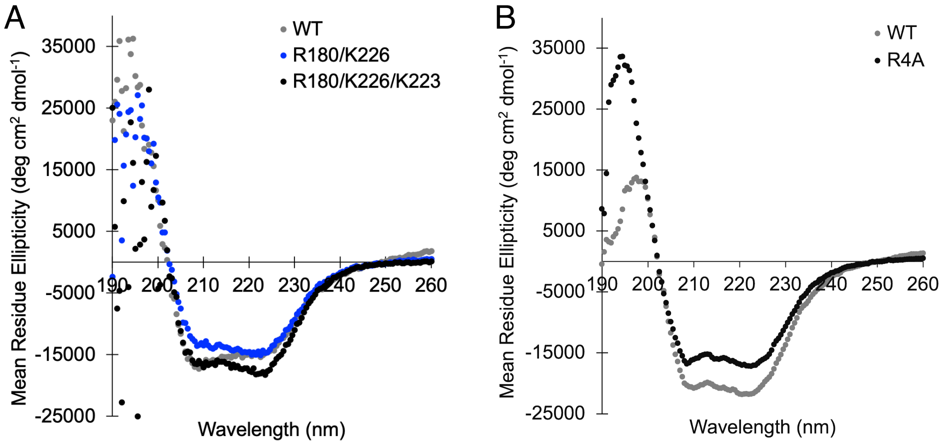


**Figure S4: Mutating the substrate binding groove does not disrupt kinase domain structure.** Circular dichroism in the far-UV region of MBP-Hhp1ΔC (A) and MBP-CK1εΔC (B) wildtype and mutant proteins.

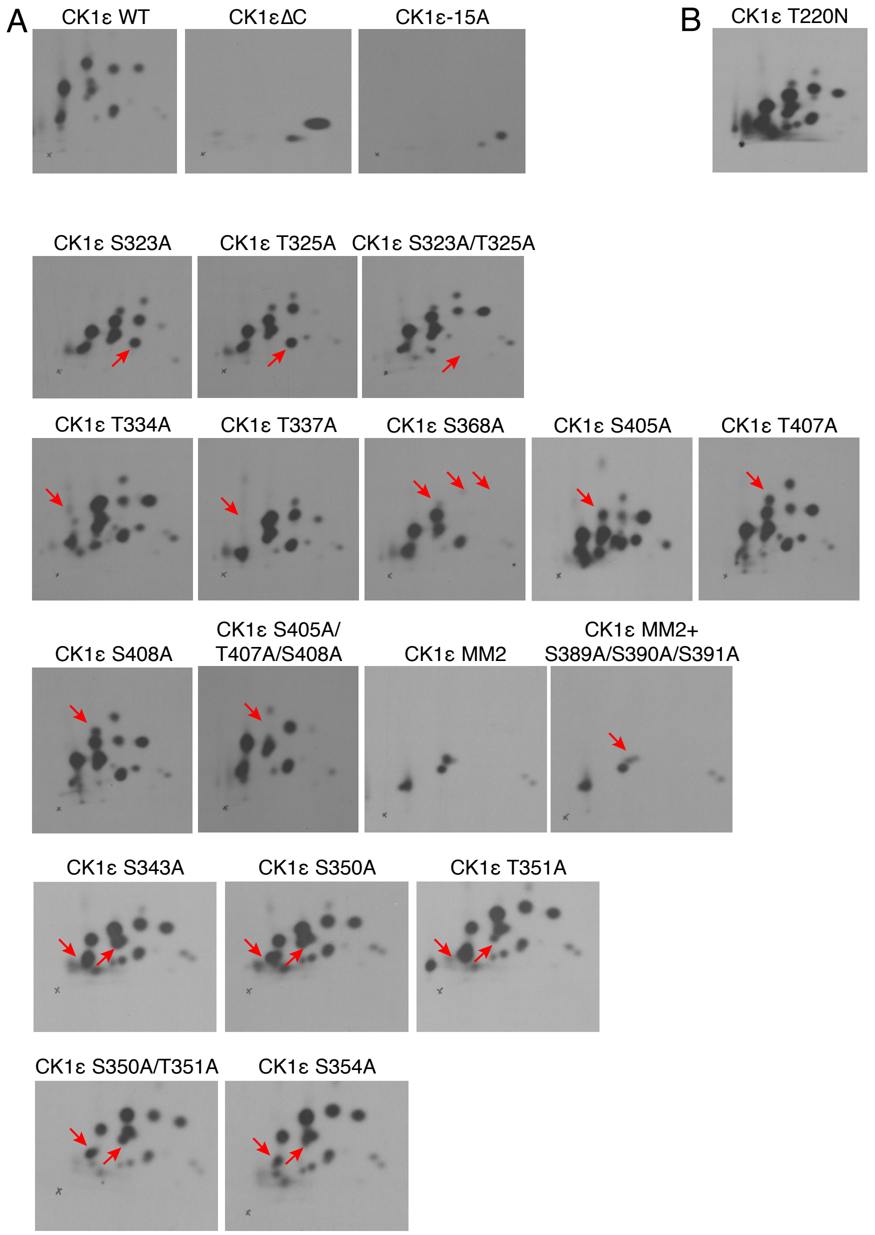


Supplemental Figure 5

**Figure S5: Identification of CK1ε autophosphorylation sites.** (A) The indicated CK1ε mutants were treated with lambda phosphatase, then incubated with γ-[$^{32}$P]-ATP at 30°C for 30 min. Proteins were digested with trypsin, and peptides were separated by thin-layer electrophoresis and chromatography. Phosphopeptides were detected by autoradiography. Red arrows point out phosphopeptides affected by alanine substitutions. CK1ε-MM2 consists of S323A, T325A, T334A, T337A, S368A, S405A, T407A, and S408A (Gietzen and Virshup, 1999). (B) Phosphopeptide map of C-terminal autophosphorylation sites in MBP-CK1ε-T220N.

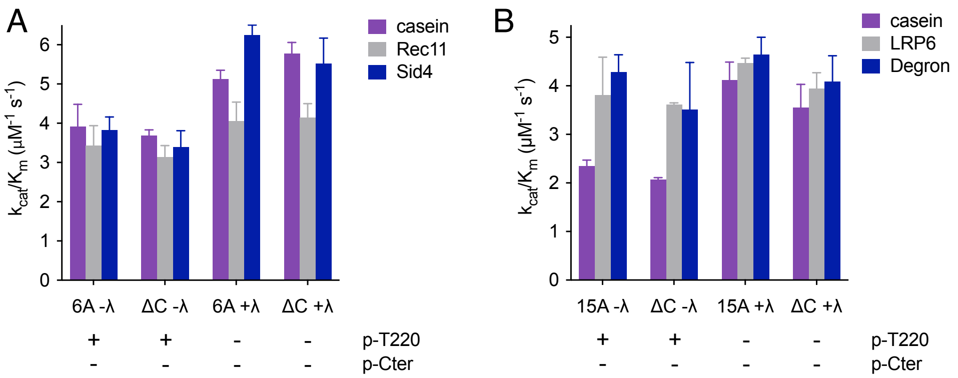


**Figure S6: Specificity footprints of Hhp1-6A and CK1ε-15A.** The indicated CK1 enzymes were treated +/- lambda phosphatase (λ), then incubated with the indicated substrates and γ-[$^{32}$P]-ATP at 30°C. Reactions were quenched at timepoints from 0-60 min, and substrate phosphorylation was measured on a phosphorimager. The initial rate was determined by the slope of the linear section of the curve, then used to calculate the $k_{cat}/K_m$ (see Methods for details). Data from three independent replicates is shown as the mean ± SD.

**Supplemental Table 1: *S. pombe* strains used in this study.**

| Strain | Genotype | Reference |
|---|---|---|
| Figure 1 | | |
| 599 | *ura4-D18 leu1-32 h-* | Lab stock |
| 14041 | *hhp1-3xFLAG:kanMX6 ura4-D18 leu1-32 ade6-M210 h-* | This study |
| 6961-2 | *hhp1-6N-3xFLAG:kanMX6 ura4-D18 leu1-32 h-* | This study |
| 7549-2 | *hhp1-K40R-3xFLAG:kanMX6 ura4-D18 leu1-32 h-* | This study |
| Supplemental Figure 2 | | |
| 599 | *ura4-D18 leu1-32 h-* | Lab stock |
| 6415 | *hhp1Δ::ura4$^{+}$ ura4-D18 leu1-32 h-* | Bimbo et al. 2005 |
| 2780-2 | *hhp1-6A ura4-D18 leu1-32 h-* | This study |
| 5592-2 | *hhp1-6N ura4-D18 leu1-32 h-* | This study |
| 3415-2 | *hhp1-6E ura4-D18 leu1-32 h-* | This study |
| * | *ade6-M210/ade6-M216 ura4-D18/ura4-D18 leu1-32/leu1-32 h-/h+* | This study |
| * | *hhp1$^{+}$/hhp1-6A ade6-M216/ade6-M210 ura4-D18/ura4-D18 leu1-32/leu1-32 h-/h+* | This study |
| * | *hhp1-6A/hhp1-6A ade6-M210/ade6-M216 ura4-D18/ura4-D18 leu1-32/leu1-32 h-/h+* | This study |
| * | *hhp1$^{+}$/hhp1-6E ade6-M216/ade6-M210 ura4-D18/ura4-D18 leu1-32/leu1-32 h-/h+* | This study |
| * | *hhp1-6E/hhp1-6E ade6-M210/ade6-M216 ura4-D18/ura4-D18 leu1-32/leu1-32 h-/h+* | This study |
| 16788 | *hhp1-mNeonGreen:kanMX6 ura4-D18 leu1-32 ade6-M210 h-* | Elmore et al. 2018 |
| 5466-2 | *hhp1-6N-mNeonGreen:kanMX6 ura4-D18 leu1-32 h-* | This study |

*Diploids were made fresh immediately prior to each experiment.
